# Synergistic Role of Amino Acids in Enhancing mTOR Activation Through Lysosome Positioning

**DOI:** 10.1101/2024.10.12.618047

**Authors:** Oralia M. Kolaczkowski, Baley A. Goodson, Valeria Montenegro Vazquez, Jingyue Jia, Aadil Qadir Bhat, Tae-Hyung Kim, Jing Pu

## Abstract

Lysosome positioning, or lysosome cellular distribution, is critical for lysosomal functions in response to both extracellular and intracellular cues. Amino acids, as essential nutrients, have been shown to promote lysosome movement toward the cell periphery. Peripheral lysosomes are involved in processes such as lysosomal exocytosis, cell migration, and metabolic signaling—functions that are particularly important for cancer cell motility and growth. However, the specific types of amino acids that regulate lysosome positioning, their underlying mechanisms, and their connection to amino acid-regulated metabolic signaling remain poorly understood. In this study, we developed a high-content imaging system for unbiased, quantitative analysis of lysosome positioning. We examined the 15 amino acids present in cell culture media and found that 10 promoted lysosome redistribution toward the cell periphery to varying extents, with aromatic amino acids showing the strongest effect. This redistribution was mediated by promoting outward transport through SLC38A9-BORC-kinesin 1/3 axis and simultaneously reducing inward transport via inhibiting the recruitment of Rab7 and JIP4 onto lysosomes. When examining the effects of amino acids on mTOR activation—a central regulator of cell metabolism—we found that the amino acids most strongly promoting lysosome dispersal, such as phenylalanine, did not activate mTOR on their own. However, combining phenylalanine with arginine, which activates mTOR without affecting lysosome positioning, synergistically enhanced mTOR activity. This synergy was lost when lysosomes failed to localize to the cell periphery, as observed in kinesin 1/3 knockout (KO) cells. Furthermore, breast cancer cells exhibited heightened sensitivity to phenylalanine-induced lysosome dispersal compared to noncancerous breast cells. Inhibition of LAT1, the amino acid transporter responsible for phenylalanine uptake, reduced peripheral lysosomes and impaired cancer cell migration and proliferation, highlighting the importance of lysosome positioning in these coordinated cellular activities. In summary, amino acid-regulated lysosome positioning and mTOR signaling depend on distinct sets of amino acids. Combining lysosome-dispersing amino acids with mTOR-activating amino acids synergistically enhances mTOR activation, which may be particularly relevant in cancer cells.

## INTRODUCTION

Lysosomes are dynamic cellular organelles with diverse functions beyond macromolecule digestion. They play pivotal roles in autophagy, pathogen elimination, metabolic signaling, gene regulation, cell adhesion and migration, as well as cell death processes^1, 2^. Many of these functions depend on lysosome dynamics, which are influenced by their movement and positioning within the cell. Lysosomes move along microtubule tracks to various cellular regions or remain stationary by interacting with other organelles^3, 4^ or actin filaments^5^. This results what is known as lysosome positioning, which is crucial for lysosomal and cell functions^1, 2, 6, 7^.

Lysosomes move in two directions: anterograde transport, which moves lysosomes from the microtubule organizing center (MTOC) near the nucleus to the cell periphery, and retrograde transport, which returns lysosomes toward the MTOC. Anterograde transport is driven by motor proteins kinesin 1 and kinesin 3^8-10^, while retrograde transport relies on the dynein-dynactin motor complex^11, 12^. Lysosome movement is mediated by various lysosomal membrane proteins and adaptor proteins, which form protein-protein interaction chains to couple lysosomes with motor proteins^1, 2, 6, 7^. Another major class of regulators are lysosomal small GTPases, such as ARL8 and Rab7, which regulate both anterograde and retrograde transport. ARL8, for example, links lysosome membrane associated BORC complex to kinesin 3 or, alternatively, to kinesin 1 via its effector SKIP^10, 13-15^. It also connects lysosomes to the dynein-dynactin complex via RUFY3 and RUFY4^16, 17^. Rab7 mediates lysosome attachment to kinesin 1 through FYCO1^18, 19^ and to dynein–dynactin via RILP^20, 21^. Interestingly, ARL8 and Rab7 share several interacting proteins, such as PLEKHM1^22^ and TBC1D15^23^, which provide potential mechanisms for crosstalk between the two GTPases. However, how ARL8 and Rab7 selectively switch between their effectors remains unclear. It is possible that additional regulatory mechanisms influence their preference for specific effectors in response to different cellular cues.

Lysosome positioning responds to various extracellular and intracellular cues, such as nutrient availability, pathogen infection, pH change, oxidative stress, protein aggregation, and oncogenesis^1, 2^. Changes in lysosome positioning under different regulatory inputs spatiotemporally influence lysosomal functions. For example, the removal of amino acids and serum from cell culture media induces autophagy and causes lysosome re-localization to the juxtanuclear area ^4, 24^, where autophagosomes accumulate. This juxtanuclear clustering of lysosomes enhances the rate of lysosome-autophagosome fusion, supporting normal autophagic flux^24, 25^. In contrast, when amino acids and serum are present, lysosomes move toward the cell periphery, where mTOR—a central regulator of cell metabolism—is recruited to lysosomes and activated through the mTOR-contained protein complexes mTORC1^24^ and mTORC2^26^. The relationship between lysosomal positioning and mTOR activity highlights the role of lysosomes as dynamic signaling hubs, facilitating efficient signal transduction. Despite the significance of this interplay, the precise mechanisms connecting lysosome positioning to mTOR activity remain unclear. One proposed mechanism suggests that the pathways regulating mTOR and lysosomal positioning converge at the lysosomal amino acid transceptor SLC38A9 and mTOR regulatory complex Ragulator^27, 28^. Additionally, the production of phosphatidylinositol 3-phosphate (PtdIns3P), a factor known to activate mTOR and disperse lysosomes, may also link these processes^19^.

Amino acids are the building blocks of protein synthesis and play key roles as intermediates in various metabolic pathways. Beyond activating mTORC1, amino acids regulate critical cellular processes and display specific roles depending on cell type^29^. For instance, metabolites of aromatic amino acids serve as precursors for neurotransmitters in neurons, whereas in cancer cells, they contribute to carcinogenesis and immunosuppression^30^. These distinct functions underscore the complexity of amino acid-mediated regulation and emphasize the need to study individual amino acids in specific contexts. Given the importance of mTOR signaling and lysosome positioning in cancer progression^1, 31-33^, it is crucial to understand how amino acids influence these processes in mechanisms and functions. In line with this, the large neutral amino acid transporter 1 (LAT1, or SLC7A5), a known activator of mTORC1^34, 35^, has been implicated in cancer progression and represents a promising target for therapeutic intervention^36, 37^.

Distinct from mTOR signaling studies, the methods for analyzing lysosome positioning predominantly rely on microscopy imaging and subsequent image analysis^38^. While electron microscopy can identify lysosomes without labeling, fluorescence microscopy, which is more commonly used, typically requires labeling with lysosome-specific dyes, cargo molecules, or antibodies against lysosomal marker proteins^38^. Common lysosome markers include LAMP1 and LAMP2, which are also found on organelles that share features with lysosomes, such as late endosomes and autolysosomes. Given the similarities among these organelles, we refer to all LAMP1/2-positive organelles collectively as “lysosomes” in this study. To quantitatively assess lysosome positioning, two strategies are frequently employed: one involves quantifying lysosome numbers within defined concentric regions or "shells" generated by shrinking the cell boundary at fixed intervals, and the other measures the distance between individual lysosomes and a reference object, such as the nucleus or MTOC^38, 39^. While automated image analysis software can reduce bias and expedite quantification, microscopy images are still often collected manually, which introduces the potential for unintentional bias and is time-consuming. Additionally, due to the limitations of these methods, only a small number of cells from limited experimental groups are typically analyzed, making them unsuitable for high-throughput or screening applications.

In this study, we developed a method using an automated high-content imaging system to quantitatively analyze lysosome positioning, ensuring unbiased imaging and data analysis. With this approach, we identified that 10 out of 15 amino acids in cell culture media were able to disperse lysosomes through promoting SLC38A9-BORC-kinesin-mediated anterograde transport and inhibiting Rab7- and SLC38A9-JIP4-mediated retrograde transport. Aromatic amino acids were the most efficient at driving lysosomes toward the cell periphery. This peripheral lysosome localization synergized with mTOR-activating amino acids, such as arginine, to enhance mTOR activation. Furthermore, we found that phenylalanine dispersed lysosomes more rapidly and strongly in breast cancer cells compared to noncancerous cells. Inhibition of LAT1, which mediates the uptake of large neutral amino acids like phenylalanine, impaired lysosome peripheral distribution and cell migration and proliferation. We conclude that amino acids functionally separate in their regulation of lysosome positioning and mTOR activation. Importantly, the localization of lysosomes to the cell periphery facilitated mTOR activation when different types of amino acids were present simultaneously. This synergy may play an important role in cancer cell motility and proliferation.

## RESULTS

### Identification of amino acids that regulate lysosome positioning

By comparing the effects of short-term (1 h) depletion of different nutrient components in HeLa cells, we demonstrated that amino acid depletion resulted in the clustering of lysosomes in the juxtanuclear region, a distribution pattern similar to that observed during complete starvation (Fig. 1A). This clustering effect was more pronounced than that seen with glucose or serum starvation (Fig. 1A), indicating that lysosome positioning is particularly sensitive to amino acid availability. Furthermore, we confirmed that this response is not cell-type specific by showing that both monkey COS-7 cells and human U2OS cells also exhibited lysosome clustering under amino acid-depleted conditions (Fig. S1A), suggesting that the response of lysosomes to amino acid depletion is a conserved mechanism across different cell lines.

**Fig. 1.**
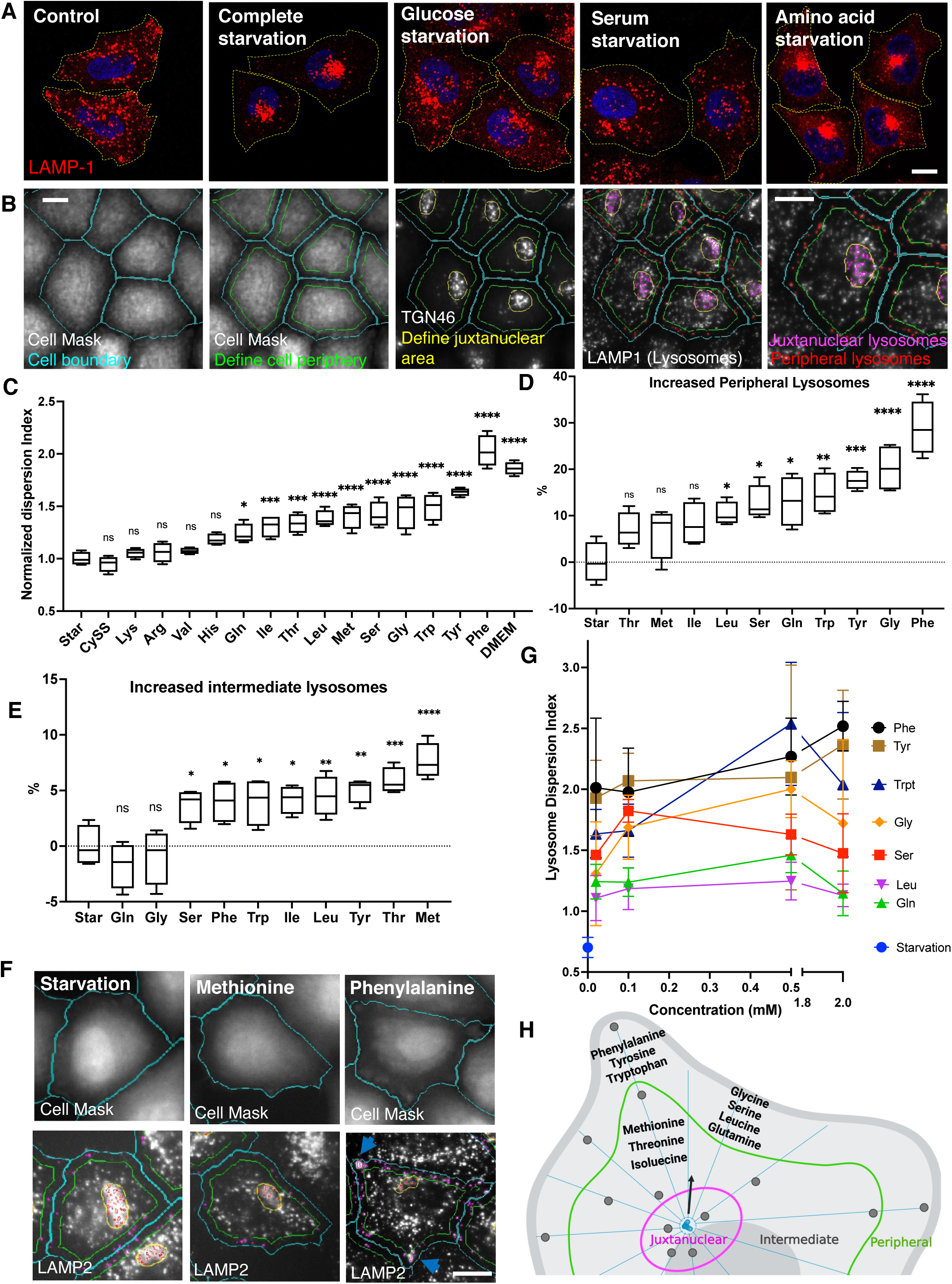
Identification of lysosome-dispersing amino acids by high-content imaging analysis. **A.** HeLa cells were starved in different nutrient-depleted media for 1 h and fixed immediately for immunostaining with an antibody against LAMP1. Fluorescence confocal microscopy was performed to visualize lysosome cellular distribution. Yellow doted lines were added to indicate the cell boundaries. **B.** An image example of high-content imaging analysis by CellInsight for lysosome positioning. CellMask was applied to identify cell boundaries, and a shell was made by shrinking cell boundary inward with a 15-micron gap (cell peripheral area). The signal of immunostaining of TGN46 was smoothened and identified as a circle to define juxtanuclear area. Lysosomes were immunostained with a LAMP1 antibody. All the masks were automatedly generated by the software. **C.** Cells were starved in amino acid- and serum-free media for 1 h and then refed with 2 mM individual amino acid or DMEM (no serum) for 30 min, followed by the lysosome positioning analysis as described in B. Lysosome Dispersion Index is defined as the ratio of cell peripheral to juxtanuclear lysosome fluorescence intensity and normalized to the average value of starvation group. The percentages of increased cell peripheral (**D**) or intermediate (**E**) lysosomes out of total lysosomes were shown for the 10 amino acids that increased the Lysosome Dispersion Index in C. Example images obtain by CellInsight were shown in **F**. Note that phenylalanine accumulated lysosomes at cell protrusions pointed by arrows. **G.** Cells were starved as in C and refed with the indicated 7 amino acids at the concentrations of 0.02, 0.1, 0.5, and 2 mM. **H.** A schematic cartoon showing different types of amino acids disperse lysosomes to different cell areas. Floating bar graphs are presented as min-max and mean ± SD. Points in the curve graph are presented as mean ± SD. *p* values were determined using One-way ANOVA test. *, *p*<0.05, **, *p*<0.001, ***, *p*<0.001, ****, *p*<0.0001, n.s., not significant (vs. Starvation). Scale bars, 5 μm.

To identify which amino acids regulate lysosome positioning, we established an image-based high-content assay using the automated CellInsight microscopy platform to ensure unbiased image collection and analysis. Cells were stained with a cytoplasmic dye and an antibody against the trans-Golgi network (TGN) marker protein TGN46, which allowed for the clear identification of cell boundaries and the MTOC area, defining the peripheral and juxtanuclear regions, respectively (Fig. 1B). Lysosomes were immunostained with an antibody against LAMP1, and their fluorescence intensities were quantified within the defined cell sections (Fig. 1B). To quantify the extent of lysosome distribution, we applied a metric termed the Lysosome Dispersion Index, which represents the ratio of peripheral lysosomes to juxtanuclear lysosomes. This index provides a quantitative measure of lysosome dispersal from the juxtanuclear area to the cell periphery, facilitating a clear assessment of lysosome positioning.

Next, we examined 15 amino acids present in DMEM and in human blood, including 9 essential amino acids—histidine, isoleucine, leucine, lysine, methionine, phenylalanine, threonine, tryptophan, and valine—and 6 non-essential amino acids—arginine, cystine, glutamine, tyrosine, glycine, and serine (Table 1). To compare their ability to disperse lysosomes, we refed amino acid- and serum-deprived cells with 2 mM of each amino acid individually for 30 minutes. The results indicated that 5 amino acids—cystine, lysine, arginine, valine, and histidine—had limited or no effect on lysosome positioning. In contrast, the other 10 amino acids promoted lysosome redistribution toward the cell periphery, as evidenced by significantly increased Lysosome Dispersion Indexes (Fig. 1C). Among these, aromatic amino acids— phenylalanine, tyrosine, and tryptophan—exhibited the highest dispersion indexes (Fig. 1C). Notably, phenylalanine not only dispersed lysosomes but also accumulated a population of lysosomes at cell protrusions (Figs. 1D,F, arrows). In contrast, methionine, which produced a lower dispersion index, relocalized lysosomes to the “intermediate area” (Fig. 1F), defined as the cellular region between the periphery and juxtanuclear areas (Fig. 1H). To further assess the differences in lysosome distribution, we quantified the number of lysosomes in the peripheral and intermediate areas. Methionine, threonine, and isoleucine did not significantly increase peripheral lysosomes but instead relocated lysosomes to the intermediate area (Figs. 1D,E). Conversely, the other 7 amino acids—including aromatic amino acids, glycine, glutamine, serine, and leucine—promoted lysosomes moving to the cell periphery (Fig. 1D).

Except for glutamine, the concentrations of other amino acids in DMEM are lower than the 2 mM applied in the experiments above, and in human blood, these levels are even lower (Tab. S1). For instance, phenylalanine concentrations are 0.4 mM in DMEM and range from 0.035 to 0.085 mM in adult blood. To determine if amino acids can influence lysosome positioning at their physiological concentrations, we examined 7 amino acids known to disperse lysosomes to the cell periphery at concentrations of 0.02, 0.1, 0.5, and 2 mM. Compared to starvation, all 7 amino acids increased the Lysosome Dispersion Index at 0.02 mM (Fig. 1G). Statistical analysis revealed that at this concentration, phenylalanine, tyrosine, tryptophan, and serine significantly increased the dispersion index (p=0.0004, 0.0007, 0.0081, and 0.0335, respectively, compared to starvation). At 0.1 mM, glycine and glutamine also showed significant increases (p=0.0002 and 0.0469). At 0.5 mM, leucine significantly enhanced the index (p=0.0023). These results suggest that amino acids can effectively regulate lysosome positioning even at physiological concentrations.

In addition to lysosomes, we also examined the positioning of other cellular organelles and found that EEA1- or Rab5-positive early endosomes clustered at the MTOC in response to starvation (Fig. S1B). This clustering was also observed during long-term serum depletion^40^. Interestingly, refeeding with phenylalanine reversed this redistribution, similar to the observed effects on lysosomes (Fig. S1B). In contrast, other organelles, including the endoplasmic reticulum (ER), mitochondria, lipid droplets, and autophagosomes, did not exhibit a change in their positioning during amino acid depletion and refeeding (Fig. S1B).

Altogether, our findings demonstrate that amino acids have distinct roles in regulating lysosome positioning (Fig. 1H). Notably, aromatic amino acids are particularly effective in promoting the distribution of lysosomes towards the cell periphery, an effect that appears to be specific to the endolysosomal vesicles. In this study, we primarily focused on the regulations to lysosomes.

### Structural basis of aromatic amino acids in regulating lysosome positioning

The structural similarities among aromatic amino acids suggest that the aromatic ring plays a crucial role in their regulatory mechanisms. To investigate whether this ring structure is sufficient to elicit the observed regulatory effects, we examined various phenylalanine metabolites and a derivative that retain the aromatic ring but differ in other structural components (Fig. 2A). For instance, p-Fluoro-dl-phenylalanine has the hydrogen at position 4 on the aromatic ring replaced by a fluoro group. This modification prevents phenylalanine hydrolase from converting the compound to tyrosine, yet our results indicate that it does not diminish the compound’s ability to promote lysosome dispersion (Fig. 2B). Among the metabolites of phenylalanine, 3-Iodo-L-tyrosine significantly increased the Lysosome Dispersion Index, while other metabolites had little or limited effect (Fig. 2C). Notably, those lacking the amino acid backbone—such as N-acetyl-L-phenylalanine, phenylpyruvate, trans-cinnamic acid, 4-hydroxyphenylpyruvic acid, phenol, and tyramine—showed minimal or no impact on lysosome dispersion despite containing an aromatic ring (Fig. 2A). These findings suggest that the aromatic ring alone is insufficient for promoting lysosome positioning, highlighting the necessity of the amino acid backbone structure. Given that not any amino acid can regulate lysosome positioning, the ability of amino acids to disperse lysosomes depends on both the backbone and the side-chain structure, highlighting the importance of their specific chemical properties.

**Fig. 2.**
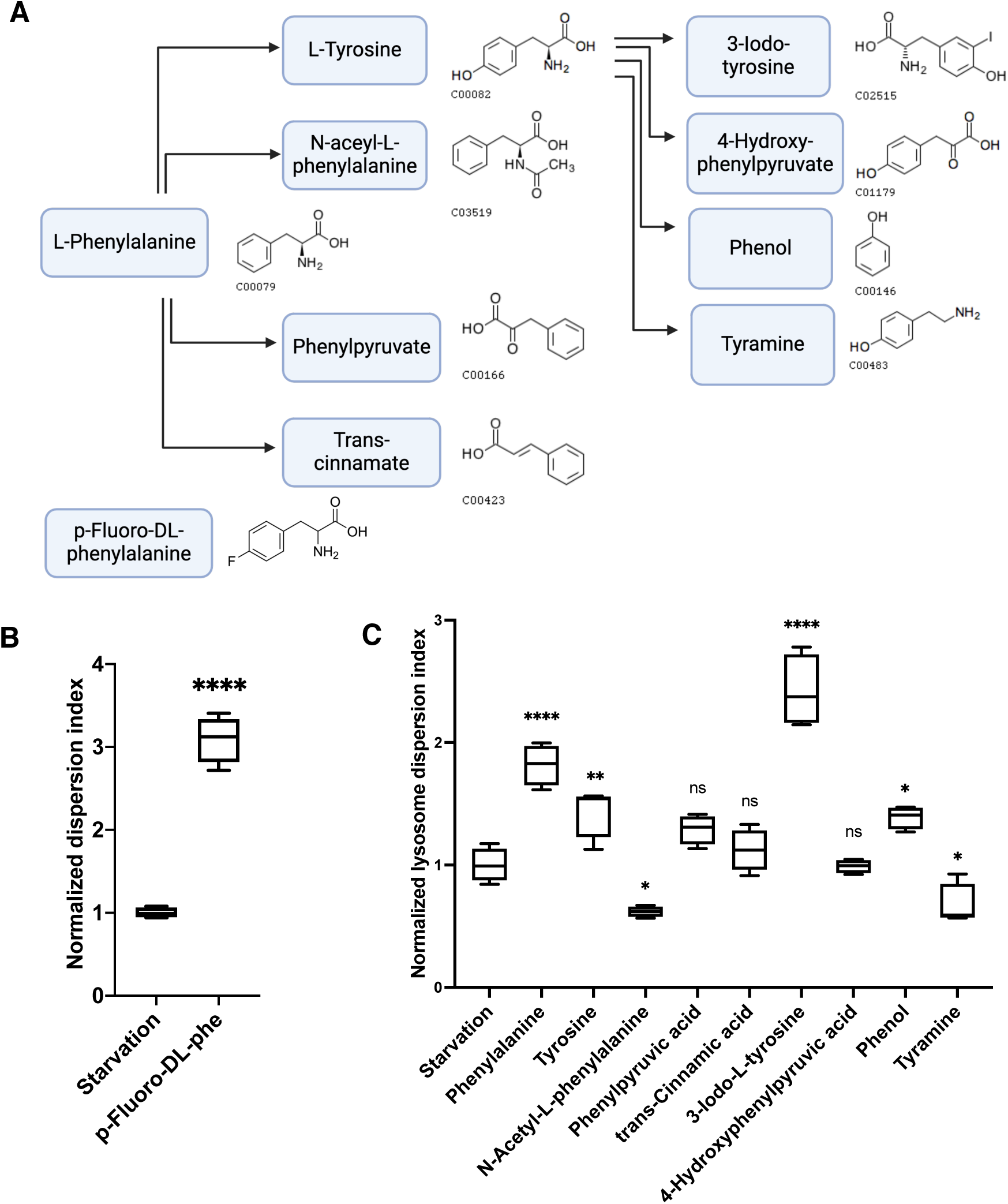
Structural basis of aromatic amino acids in dispersing lysosomes. **A.** Partial phenylalanine metabolism pathways and metabolites’ structures. The compound code and pictures of molecular structure were from KEGG Pathway or manufactures. **B. & C.** Cells were starved for 1 h and treated with phenylalanine, its metabolites, or a derivative at 2 mM for 30 min. Cells were fixed and immunostained for lysosome positioning analysis with CellInsight. Floating bar graphs are presented as min-max and mean ± SD. *p* values were determined using *t-Student’s* test or One-way ANOVA test. *, *p*<0.05, **, *p*<0.001, ****, *p*<0.0001, n.s., not significant (vs. Starvation).

### Amino acids promote lysosome anterograde transport

Previous studies have highlighted the role of SLC38A9-regulated BORC activity in mediating amino acid-induced anterograde transport^27, 28^, and kinesin 1 and 3 are the primary motor proteins for this process^10^. In alignment with these findings, knocking out (KO) of SLC38A9^28^, BORCS5^14^, or both kinesin 1 and 3^41^ resulted in a redistribution of lysosomes toward the juxtanuclear region under nutrient-rich conditions (Fig. 3A). Upon starvation, when cells were refed with the 10 amino acids that promote lysosome dispersion, two distinct outcomes were observed: 7 amino acids (tryptophan, tyrosine, phenylalanine, glutamine, leucine, serine, and glycine) exhibited reduced Lysosome Dispersion Indexes in SLC38A9-KO and BORCS5-KO cells compared to wild-type cells (Figs. 3B,C). This suggests that SLC38A9 and BORC are essential for lysosome positioning in response to these amino acids. However, the dispersion was not completely abolished in the absence of SLC38A9 or BORC, indicating the presence of alternative regulatory mechanisms (Fig. 3E). In contrast, isoleucine, threonine, and methionine did not significantly alter lysosome positioning in SLC38A9- or BORCS5-KO cells (Fig. 3D), suggesting that the SLC38A9-BORC pathway specifically responds to these three amino acids. Interestingly, deletion of kinesin 1 and 3 completely abolished lysosome dispersion in response to all ten amino acids (Figs. 3B,C,D), confirming the role of these motor proteins in lysosomal anterograde transport^10, 14^. Collectively, these results indicate that amino acids promote lysosome anterograde transport through various regulatory mechanisms that converge on kinesins 1 and 3 (Fig. 3E).

**Fig. 3.**
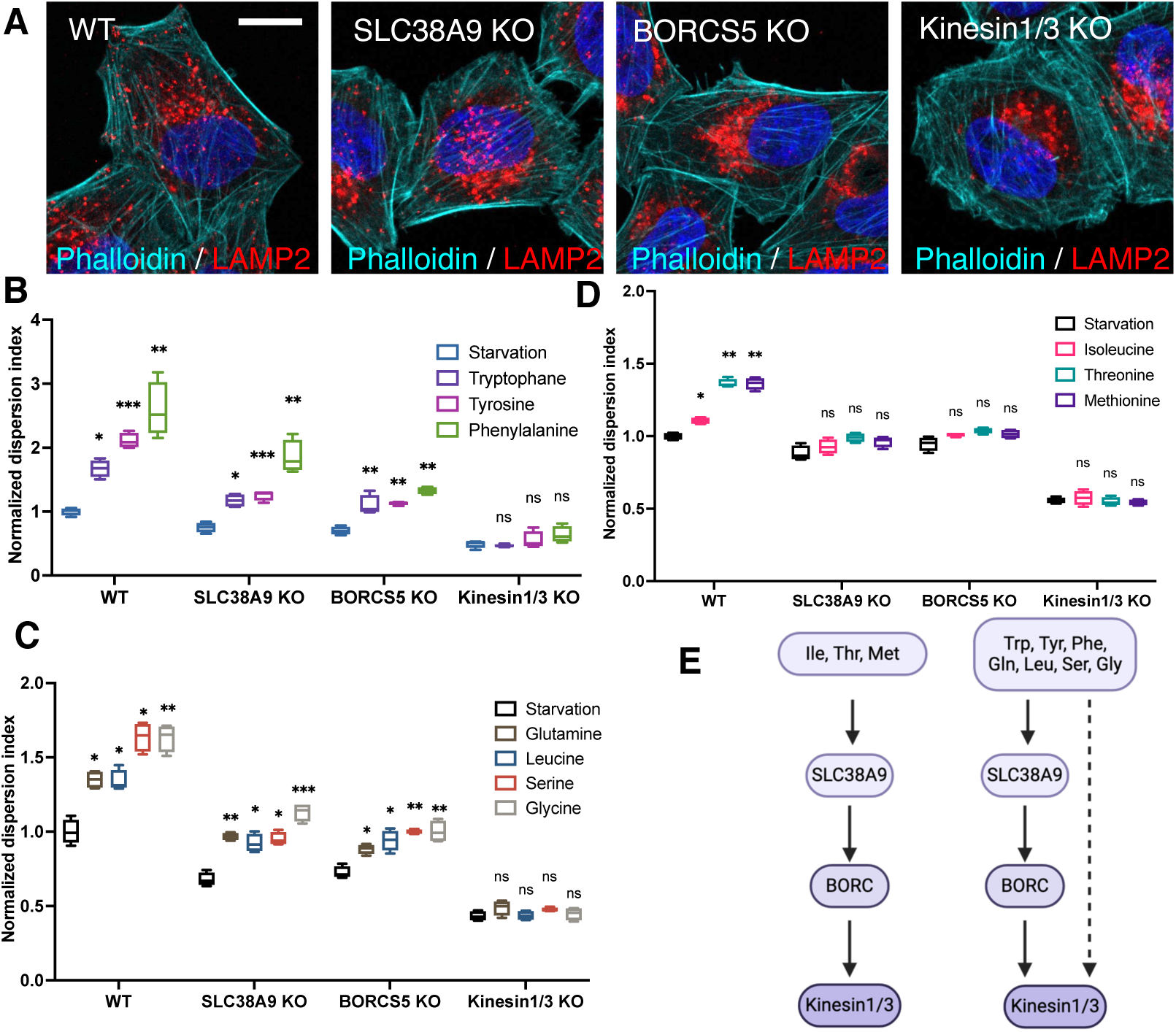
Characterization of the mechanisms regulating lysosome anterograde transport in response to amino acids. **A.** Wild-type and CRISPR-gene KO HeLa cells were immunostained with LAMP2 antibody to visualize lysosomes and stained with phalloidin (for actin) to visualize cell boundaries. **B-D,** Wildtype and CRISPR-gene KO cells were starved for 1 h and refed with indicated amino acids for 30 min. Lysosome positioning was analyzed and presented as the Lysosome Dispersion Indexes. **E**. Schematic illustration of the pathways that mediated lysosome anterograde transport in response to different amino acids. Floating bar graphs are presented as min-max and mean ± SD. *p* values were determined using One-way ANOVA test. *, *p*<0.05, **, *p*<0.001, ***, *p*<0.001, ****, *p*<0.0001, n.s., not significant (vs. Starvation in the same cell group). Scale bars, 5 μm.

### Amino acids inhibit lysosome retrograde transport

To investigate whether amino acids regulate lysosome retrograde transport, we examined three known regulators: Rab7, JIP4, and TRIML-1. These proteins couple lysosomes to the retrograde motor dynein/dynactin complex through distinct pathways^2^. We deleted Rab7 using CRISPR-Cas9 system^42^, confirmed by Sanger sequencing and immunoblotting (Fig. 4A). In Rab7-KO cells, swollen lysosomes were filled with cholesterol (Fig. S2A), consistent with reported results^43^. Rab7 mediates both anterograde and retrograde transport, and lysosomal cholesterol regulates retrograde transport^18-21, 44^, suggesting a complex role of Rab7. Our analysis of Rab7-KO cells demonstrated a general increase in Lysosome Dispersion Indexes by 1 to 3-fold under both starvation and amino acid refeeding conditions (Fig. 4B), suggesting that Rab7 primarily enhances retrograde transport in our experimental conditions.

**Fig. 4.**
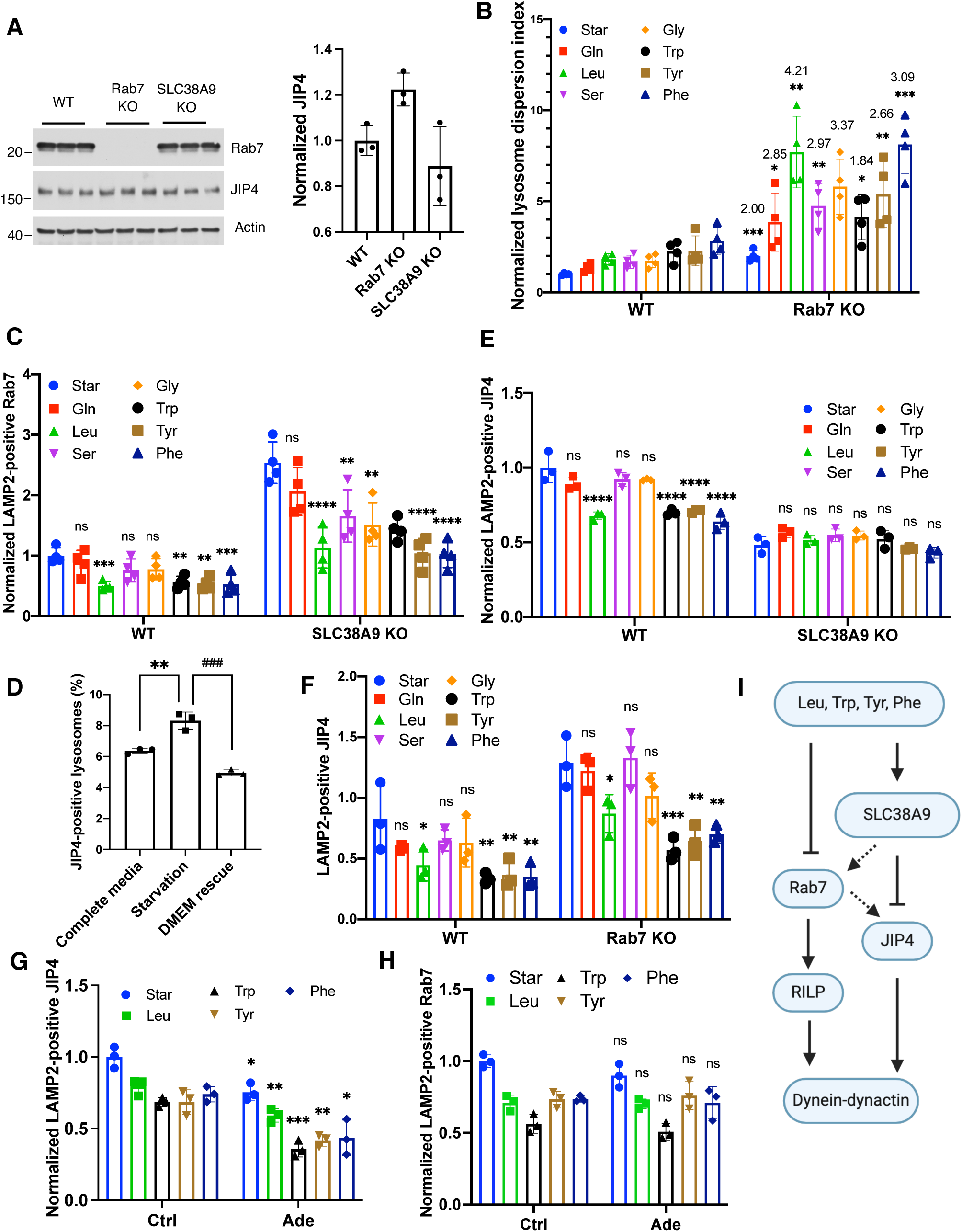
Characterization of the mechanisms regulating lysosome retrograde transport in response to amino acids. **A.** Immunoblotting with the indicated antibodies for wildtype (WT), Rab7-KO, and SLC38A9-KO cells. Three replicate samples were examined and quantified using FIJI. **B-H,** WT and indicated KO cells were starved for 1 h and refed with indicated amino acids for 30 min. When the p38 inhibitor Adezmapimod (Ade) was applied, cells were starved and refed along with the drug at 10 µM. Cells were fixed and immunostained with LAMP2, TGN46, and Rab7 or JIP4 antibodies. Lysosome positioning was analyzed and presented as lysosome dispersing indexes. LAMP2-positive Rab7 or JIP4 was identified and quantified using the “colocalization” function in the software of CellInsight. Numbers on the bars represent the fold of Rab7-KO group vs. WT group. **I**. Schematic illustration of the pathways that mediated lysosome retrograde transport in response to amino acids. Dotted arrows present potential regulation at gene expression-levels. Bar graphs are presented as mean ± SD. *p* values were determined using *t-Student’s* test or one-way ANOVA test. *, *p*<0.05, **, *p*<0.001, ***, *p*<0.001, ****, *p*<0.0001, n.s., not significant (in B, G, and H, vs. WT; in C, E, and F, vs. Starvation in the same cell group).

We further assessed lysosomal-localized Rab7 and found that leucine and aromatic amino acids reduced the number of LAMP2-positive Rab7 puncta (Fig. 4C), indicating that these four amino acids effectively repel Rab7 from lysosomes. Given that these amino acids promote lysosome dispersion (Fig. 1C), the reduction of lysosomal Rab7 suggests that the primary role of Rab7 is coupling lysosomes to dynein-dynactin (Fig. 4C). In addition, deletion of SLC38A9 resulted in an increased number of LAMP2-positive Rab7 puncta (Fig. 4C) without altering the overall protein level (Fig. 4A). Importantly, the amino acid-induced disassociation of Rab7 from lysosomes was still observed in SLC38A9-KO cells (Fig. 1C), indicating that while SLC38A9 deletion may constitutionally enhance Rab7 association with lysosomes (possibly as a cellular adaptation), it does not affect the acute regulation by amino acids.

Next, we investigated JIP4 (also known as SPAG9), which couples lysosome membrane protein TMEM55B to dynein-dynactin^45^. We found that starvation slightly but significantly increased JIP4 levels at lysosomes, which was abolished by amino acid refeeding (DMEM) (Fig. 4D). This suggests that JIP4 dynamically associates with lysosomes in response to amino acid availability and plays a role in lysosome retrograde transport. Notably, leucine and aromatic amino acids significantly reduced LAMP2-positive JIP4 puncta (Fig. 4E), and this reduction was blocked by SLC38A9 deletion (Fig. 4E) but not by Rab7 deletion (Fig. 4F). These findings imply that SLC38A9 regulates JIP4 in response to amino acids, paralleling the Rab7 pathway. Further evidence came from using a p38 MAPK inhibitor to inhibit JIP4^45, 46^. We observed a reduction in lysosomal JIP4 levels (Fig. 4G) without affecting lysosomal Rab7 levels (Fig. 4H). Additionally, in Rab7-KO cells, we noted an overall increase of lysosomal JIP4 puncta (Fig. 4F), likely due to increased expression levels of JIP4 (Fig. 4A). This suggests potential crosstalk between the Rab7 and JIP4 pathways at the level of gene expression but not signaling. Altogether, our results demonstrate that leucine and aromatic amino acids weakened lysosome retrograde transport by concurrently inhibiting both Rab7- and SLC38A9-JIP4-mediated pathways (Fig. 4I).

TRPML1, a lysosomal calcium channel, is activated by starvation and recruits downstream effectors that bind to dynein-dynactin to enhance retrograde transport^47^. Using the TRPML1 agonist ML-SA5, we found that activating TRPML1 during amino acid presence did not diminish lysosome dispersal (Fig. S2B), suggesting that TRPML1 is not involved in amino acid-regulated lysosome positioning.

### Role of lysosome positioning in synergistic mTOR activation

An important cellular response to amino acid availability is the activation or inhibition of mTOR signaling, which regulates cell growth and metabolism. Previous studies have reported differential regulation of mTOR by various amino acids^48, 49^. Using phosphorylated S6 kinase (p-S6K) as an indicator of mTOR activity, our results demonstrate that glutamine, glycine, arginine, histidine, leucine, isoleucine, serine, and threonine can activate mTOR to varying extents (Figs. 5A,B). Notably, aromatic amino acids did not stimulate mTOR, which is consistent with previous reports^49^. To illustrate the correlation between amino acid-regulated lysosome positioning and mTOR activation, we plotted the quantitative results of the Lysosome Dispersion Index (Fig. 1C) against mTOR activity (Fig. 5A). As shown in Figure 5C, cysteine, lysine, and valine neither dispersed lysosomes nor activated mTOR. Arginine activated mTOR but did not alter lysosome positioning. Conversely, methionine and the aromatic amino acids dispersed lysosomes without activating mTOR. Glutamine, glycine, isoleucine, leucine, serine, and threonine promoted both processes to varying degrees.

**Fig. 5.**
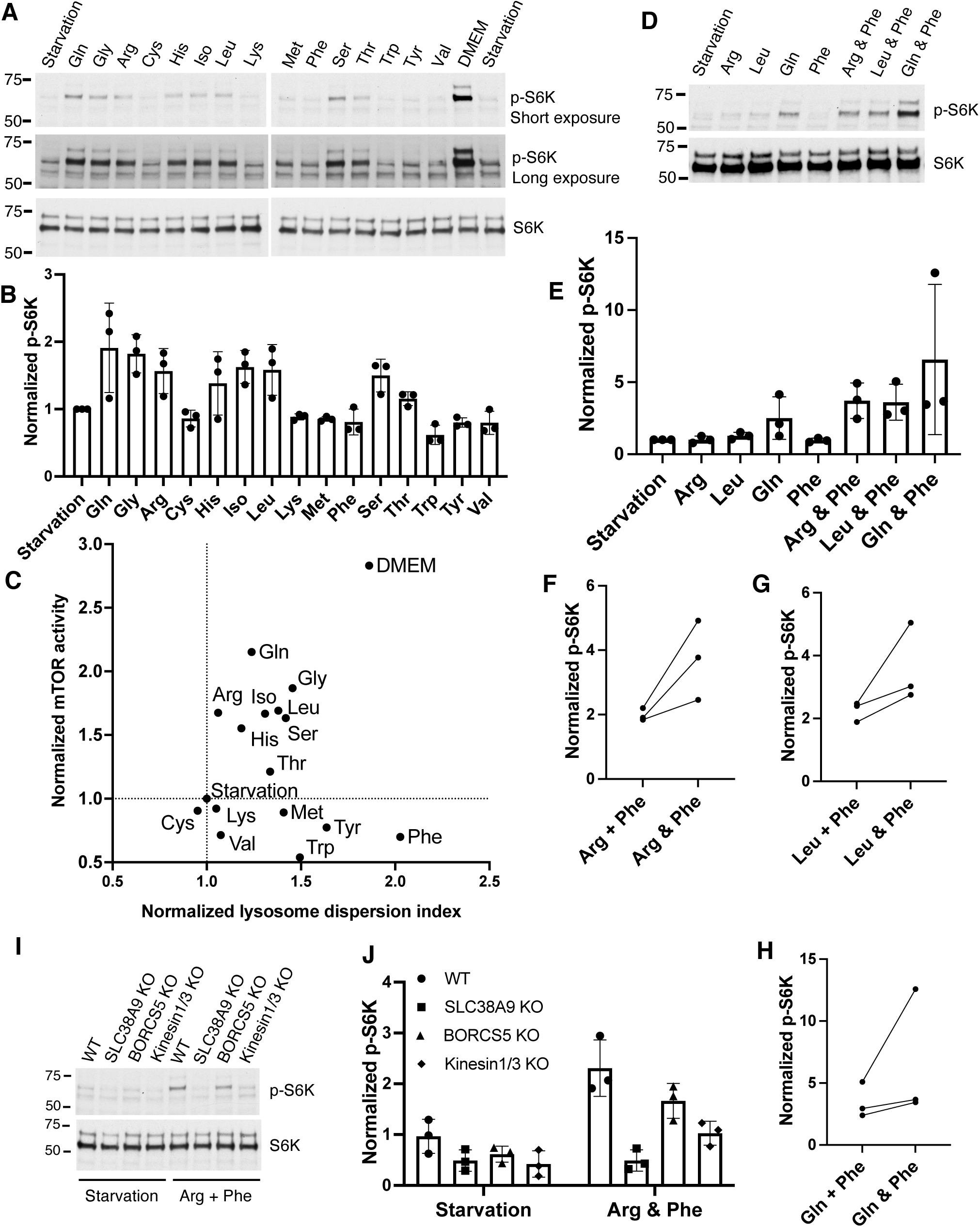
Amino acid regulation of mTOR activation. **A & B,** HeLa cells were starved and then refed with 2 mM amino acids or DMEM as indicated, lysed directly by adding 2X SDS sample buffer, and subjected to immunoblotting with the antibodies indicated. The ratio of phosphorylated S6K (p-S6K) to total S6K in each treatment was normalized to the value in starvation group. The mean± SD was plotted from 3 independent experiments. **C.** Normalized Lysosome Dispersion Indexes from Fig. 1C and mTOR activity from Fig. 5B were plotted on a coordination plane, with the X-axis representing lysosome dispersion and the Y-axis representing mTOR activity. **D & E,** Cells were treated as described in A with the indicated amino acids or amino acid combinations. Total proteins were extracted and subjected to immunoblotting. The ratio of p-S6K to total S6K for each treatment was normalized to the values in the starvation group. Data represent the mean ± SD from three independent experiments. **F-H,** The normalized values of p-S6K/S6K from individual amino acid treatments in E were summed (labeled as “+”) and compared to the corresponding values from the combination of the two amino acids (labeled as “&”). Paired values from three independent experiments were shown with lines connecting the data points. **I & J,** Wild-type (WT) and indicated gene-KO cells were treated with combined arginine and phenylalanine, and cell lysates were subjected to immunoblotting as in A. The normalized values of p-S6K/S6K from 3 independent experiments were presented as mean± SD.

Given that amino acids are mixed present in cell culture media and under physiological conditions, we explored whether amino acid-mediated lysosome positioning can regulate mTOR signaling. We specifically focused on combining two amino acids with distinct functions: arginine, which mediates mTOR activation, and phenylalanine, which promotes lysosome dispersal. Surprisingly, while phenylalanine alone has a minimal effect on activating mTOR, the combination of phenylalanine and arginine significantly enhanced mTOR activity, surpassing the effects observed with arginine alone (Figs. 5D,E,F). To determine whether this synergistic effect is exclusive to arginine, we also tested combinations of phenylalanine with leucine and phenylalanine with glutamine. Again, the combined amino acid treatments resulted in mTOR activity levels higher than those induced by leucine or glutamine alone (Figs. 5D,E,G,H). Furthermore, to assess whether phenylalanine’s role in this synergy is unique, we tested glycine and serine, both of which also promote lysosome dispersal, in the combination with arginine. Again, we observed synergistic increases in mTOR activity (Fig. S3A). These findings highlight a synergistic effect on mTOR activation driven by specific amino acid combinations.

To investigate whether lysosome positioning contributes to the observed synergistic effect between amino acids, we treated BORCS5-KO and kinesin 1/3-KO cells with the phenylalanine-arginine mixture. BORCS5-KO cells exhibited reduced mTOR activity compared to wild-type cells, while, remarkably, the deletion of kinesin 1 and 3 substantially suppressed mTOR activation (Figs. 5I,J). Similar results were obtained when these KO cells were treated with arginine-glycine and arginine-serine combinations (Fig. S3B). These findings are particularly striking, considering that BORCS5 KO did not completely block phenylalanine-mediated lysosome dispersal, while kinesin 1/3 KO did (Fig. 3B). This suggests that the peripheral distribution of lysosomes is crucial for facilitating mTOR activation.

It is known that SLC38A9 plays a vital role in mediating amino acid signaling to mTOR^50-52^. Therefore, it is not surprising that SLC38A9 deletion completely blocked mTOR activation (Figs. 5I,J & S3B), likely independent of lysosome positioning.

### Inhibition of LAT1 suppresses breast cancer cell migration and proliferation

To investigate the role of amino acid-regulated lysosome positioning in cancer, we analyzed noncancerous human mammary epithelial cells (MCF10A) alongside human breast cancer cell lines MCF7 and MDA-MB-231. A time-course study of phenylalanine refeeding following starvation demonstrated a rapid dispersion of lysosomes within just 10 minutes in both cancerous cell lines (Fig. 6A). In contrast, noncancerous MCF10A cells exhibited a significant increase of the Lysosome Dispersion Index only at the 40-minute mark, albeit at a lower level (Fig. 6A). These observations suggest that breast cancer cells are more sensitive to amino acids regarding lysosome dispersion to the cell periphery, a critical factor for cancer cell motility and invasion^32^. Furthermore, the inhibition of the amino acid transporter LAT1 using the compound KYT0353 (also known as JPH203) in the highly metastatic breast cancer cells, MDA-MB-231, led to a decreased distribution of peripheral lysosomes (Fig. 6B) and marked reduction in cell migration as early as 2 hours of treatment (Fig. 6C). This indicates that the disruption of LAT1 activity interferes with acute cellular responses, likely occurring independently of gene expression changes. Importantly, the reduced cell migration was not attributed to impaired cell proliferation, as no significant effect on cell growth was observed until the 8-hour mark (Figs. 6D,E).

**Fig. 6.**
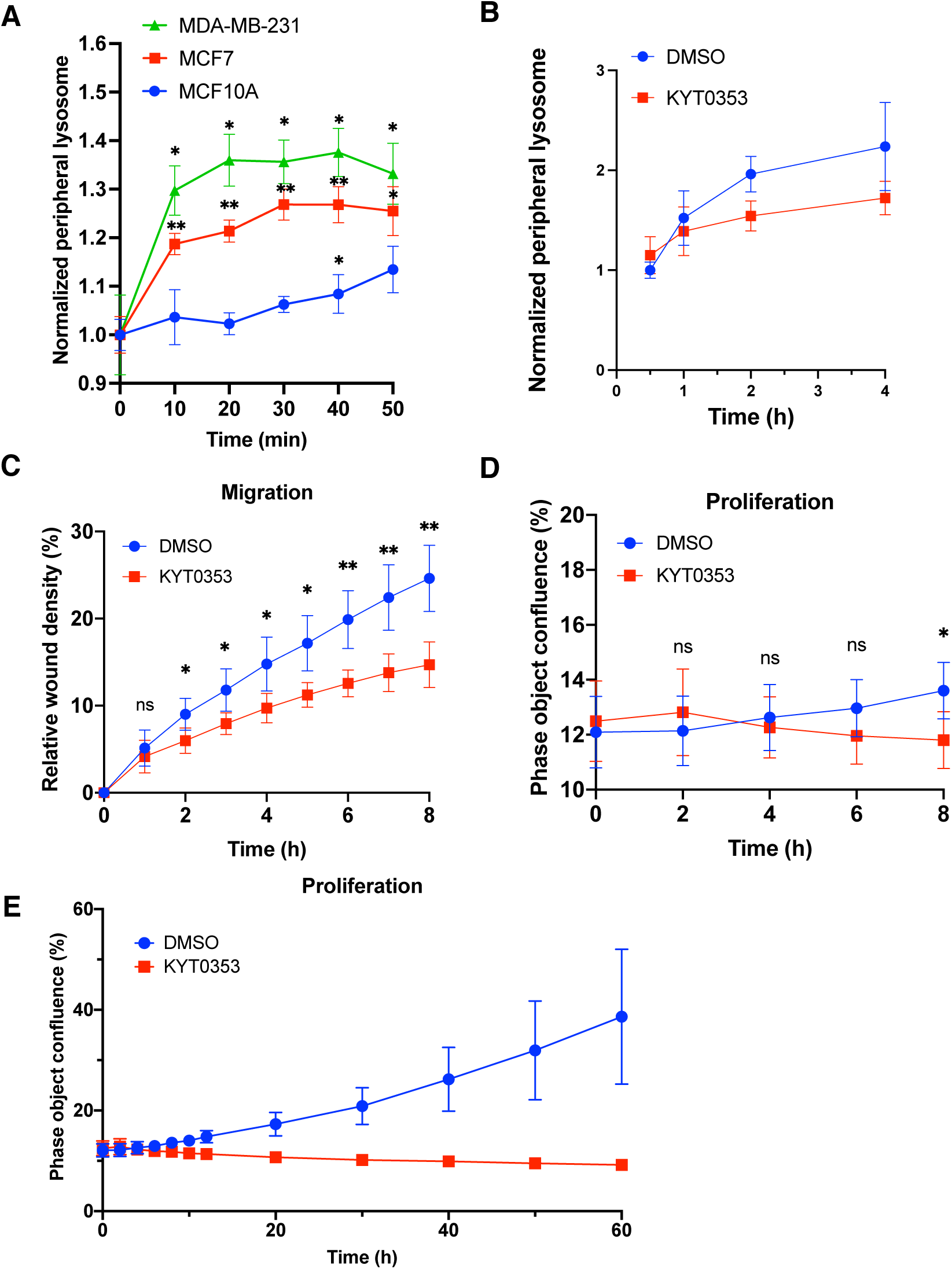
Amino acid-cell migration and proliferation in breast cancer cells. **A**. Indicated breast noncancerous (blue) and cancer cells (green and red) were starved in amino acid- and serum-free media, followed by refeeding with 2 mM phenylalanine for the indicated times. Cells were fixed, immunostaining, and analyzed by CellInsight microscope platform. Cell peripheral lysosomes were quantified from at least 1,000 cells per well, 3 wells per condition, and two independent experiments. **B.** MDA-MB-231 cells were treated with DMSO or 10 µM KYT0353 in complete media for the indicated periods and then subjected to lysosome positioning analysis as described in A. **C.** Collective cell migration assay with MDA-MB-231 cells. Cells were treated with DMSO or 10 μM KYT0353 for 60 hours. Scratch wound area was imaged hourly. Three replications were included in the assays. The data from first 8 hours were shown. **D & E.** MDA-MB-231 cells were treated with DMSO or 10 μM KYT0353 for 60 hours and imaged hourly. Cell confluency mask was quantified by using IncuCyte Zoom software. Three replications were included in the assays. The data from first 8 hours and entire treatment were shown separately. Data are presented as mean ± SD. *p* values were determined using *t-Student’s* test or one-way ANOVA test. *, *p*<0.05, **, *p*<0.001, n.s., not significant (vs. time point 0 in A. Compared between groups in B, C, and D).

## DISCUSSION

Our study identifies the amino acids that promote lysosome localization towards the cell periphery(Fig. 1C). The degree of lysosome dispersion also varies (Fig. 1C), which raises intriguing questions about the underlying mechanisms. Amino acids differ significantly in their chemical structure, and these differences likely play a key role in determining their regulatory effects on lysosomal positioning. For example, aromatic amino acids, such as phenylalanine, possess hydrophobic characteristics due to their aromatic ring structures, which allows them to interact with lipid membranes more efficiently^53^, potentially modulating membrane-associated proteins on lysosomes. In contrast, smaller, more polar amino acids may not engage lysosomal membrane proteins with the same affinity, which might explain their limited capacity to alter lysosome positioning.

We also explored how the concentration of amino acids affects lysosome positioning. Although we tested concentrations ranging from 0.02 mM to 2 mM, the changes in lysosome dispersion index were relatively modest, with no more than a 50% change (Fig. 1G). This suggests that amino acid concentration alone is not a dominant factor in regulating lysosome positioning. Instead, amino acid identity and structure likely play a more significant role.

The ability of amino acids to influence lysosome positioning may also depend on their interaction with specific lysosomal transporters and sensors. One such transporter, SLC38A9, plays a crucial role in sensing lysosomal amino acid levels and regulating outward lysosome movement by recruiting downstream effectors like kinesins^28^ (Figs. 3B,C,D) and repelling retrograde regulators such as JIP4 from the lysosomal surface (Fig. 4E). Previous studies have demonstrated that the levels of non-polar, essential amino acids like phenylalanine, leucine, isoleucine, tryptophan, and methionine in lysosomes are altered in SLC38A9 gain- and loss-of-function models^52^, aligning with our results that these amino acids can change lysosome positioning (Fig. 1C). Despite arginine binds to SLC38A9, it is not transported by SLC38A9^52^, and it does not affect lysosome positioning (Fig. 1C), suggesting that specific interactions with SLC38A9 may dictate which downstream pathways and lysosomal activities are activated. It is important to note that deletion of SLC38A9 does not entirely block the ability of certain amino acids to promote lysosome dispersal (Figs. 3B,C). This implies that additional, as-yet-unidentified mechanism might contribute to this regulation. For example, Rab7 dissociation from lysosomes, which occurs independently of SLC38A9 upon amino acid stimulation (Fig. 4C), suggests the possible existence of additional amino acid sensors that regulate lysosome positioning. However, the exact mechanisms remain elusive.

The relationship between lysosome positioning and mTOR signaling has been explored in several studies. Although mTORC1 is recruited to lysosomal surfaces for activation, its direct role in controlling lysosome positioning remains unclear. Prior research has shown that inhibiting mTOR with specific inhibitors does not significantly alter lysosome positioning^28^. In line with this, our findings indicate that boosting mTOR activity—using a combination of arginine and phenylalanine—does not further modify lysosome distribution compared to phenylalanine alone (Fig. S3C). In contrast, manipulating the molecular machinery responsible for lysosome transport has been shown to significantly influence mTOR activity^19, 24, 26^. For example, Jia and Bonifacino (2017) demonstrated that lysosomes positioned near the cell periphery, in close proximity to cell surface receptors, can enhance mTORC2 activity by facilitating access to growth factors^26^. This spatial positioning of lysosomes may be critical for nutrient uptake and activation of nutrient-sensitive pathways. Peripheral lysosomes are likely to have better access to higher local concentrations of amino acids, particularly near plasma membrane transporters. While our results suggest that amino acid concentrations do not critically influence lysosome positioning (Fig. 1G), they are essential for robust mTOR signaling^54^. Thus, lysosomes positioned at the cell periphery may localize mTORC1 signaling molecules to regions with higher amino acid concentrations, thereby enhancing mTOR activation.

## MATERIALS AND METHODS

### Cell culture

Cells used in this study, except MCF10A, were cultured in Dulbecco’s Modified Eagle’s Medium (DMEM) supplemented with 10% fetal bovine serum (FBS), 25 mM HEPES, and MycoZap Plus-CL (Lonza, Basel, Switzerland) at 37°C with 5% CO_2_. MCF10A cells were cultured in DMEM/F12 media supplemented with 5% horse serum, 20 ng/ml EGF, 0.5 mg/ml hydrocortisone, 100 ng/ml cholera toxin, 10 μg / ml insulin, 25 mM HEPES, and MycoZap Plus-CL.

### Starvation and Amino acid treatments

Cells were washed twice with amino acid-free DMEM (United State Biological, Salem, MA) supplemented with 4.5 g/L glucose, 25 mM HEPES, and MycoZap Plus-CL (AA-free medium) and incubated in the same medium for 1 h. Amino acid-2X media were prepared by adding 2-fold amino acids to AA-free medium and then mixed with the starvation media as 1:1 ratio of volume by direct addition. After 30-min treatment, cells were fixed for immunostaining or lysed to extract total proteins for immunoblotting.

### Immunostaining and confocal microscopy

Coverslips were coated with collagen solution (0.03-0.04 mg/ml collagen in 0.02 N acetic acid) for > 30 min and then washed twice with PBS before receiving cells. Cells were seeded on the coverslips a day before experiments and fixed by mixing 8% PFA in PBS and cell culture media at 1:1 volume ratio following treatments. After washing 3 times with PBS, primary antibodies were diluted in 1% BSA in PBS supplemented with 0.2% saponin and applied to cells for 1 h at 37°C or overnight at 4°C. Alexa-conjugated secondary antibodies and phalloidin (Thermo Fisher Scientific, Waltham, MA) were applied with the same saponin-contained BSA-PBS solution for 30 min at 37°C. Coverslips were mounted with DAPI-contained mounting reagents (Electron Microscopy Sciences, Hatfield, PA) and allowed to dry at 37°C for at least 1 h. Images were acquired using a Zeiss LSM800 confocal microscope equipped with the software ZEN (Zeiss, Oberkochen, Germany).

### Rab7 CRISPR-Cas9 KO

To knock out Rab7, two 20-base pair (bp) targeting sequences TAGTTTGAAGGATGACCTCT and TTGCTGAAGGTTATCATCCT were synthesized (Eurofins, Lancaster, PA) and introduced separately into the px458 plasmids containing a GFP sequence^42^ (Addgene, Cambridge, MA). HeLa cells were co-transfected with both plasmids using TransIt-LT1 (Mirus Bio, Madison, WI) following manufacture’s instruction, and GFP-positive cells were collected by cell sorting at 48 h after transfection. Transformants were kept in normal medium for another 12 days to allow single colony formation. Genomic DNA was extracted from individual colonies, and cleavage of the target sequence was tested by PCR using a pair of primers AAATAAAGCGGTCACTTCTTTGAG and ATTCCATGAAAATGCCTATCCTTA, which produced a smaller band in KO relative to WT cells. The KO was confirmed by Sanger sequencing and immunoblotting.

### High-content imaging and image analysis

Optical 96-well plates (Agilent, Santa Clara, CA) were coated with a collagen solution as previously described. Cells were seeded at 17,000 cells per well. The following day, cells were treated, fixed with 4% PFA, and subjected to immunofluorescence (IF) staining. After staining, cells were maintained in PBS for immediate imaging. Imaging and analysis were performed using the CellInsight Microscope platform (Thermo Fisher Scientific). DAPI-stained nuclei were scanned to determine focus through the programmed autofocus function. Cytoplasmic and nuclear signals were detected using HCS CellMask Near-IR Stain (Thermo Fisher Scientific) to define individual cell boundaries. Cells were excluded based on criteria such as incomplete cell boundaries at image edges, abnormal size or shape, and signs of cell death.

For lysosome positioning analysis (Fig. 1B), the cell periphery was defined as a ring region created by shrinking the cell boundary inward by 15 microns, termed ROI_A. The MTOC regions were defined by circling smoothed IF signals of TGN46, referred to as ROI_B. Lysosomes were detected as spots (2 microns in size) based on LAMP1 or LAMP2 IF signals. Lysosomes overlapping with ROI_A and ROI_B were quantified by their fluorescence intensity, representing the distribution of lysosomes in peripheral and juxtanuclear regions, respectively. The lysosomes in the intermediate area were calculated by subtracting peripheral and juxtanuclear lysosomes from the total lysosomal count.

To measure lysosomal JIP4 and Rab7, LAMP2 signals were defined as ROI_A, and the JIP4 or Rab7 signals overlapping with ROI_A were quantified for their intensity, using the software function of “colocalization”.

Each experiment included the analysis of at least 1,000 qualified cells per well, with 3-6 replicates per condition. Three or more independent experiments were performed, and the averaged data from each experiment were pooled for final analysis and plotting.

### Measuring cell migration and proliferation

To evaluate collective cell migration, a scratch wound healing assay was performed using IncuCyte Real-Time Quantitative Live Cell Analysis System (Essen BioScience, Ann Arbor, MI) as described previously^55^. Cells (6 × 10^4^) were seeded into each well of an ImageLock 96-well plate (Essen BioScience) and incubated for 24 hours to form a confluent monolayer. A scratch wound was then introduced using the WoundMaker™ (Essen BioScience). After removing cell debris with a medium wash, cells were treated with varying concentrations of KYT0353 in 100 µl of culture medium for 60 hours, with images captured every hour. Relative wound density was quantified using IncuCyte™ software (version 20158A). For cell proliferation assay, 1 x 10^4^ cells were seeded into each well of a 96-well plate and incubated for 24 hours. Following incubation, cells were treated with varying concentration of KYT0353 for 60 hours, with images captured every hour. Cell proliferation was analyzed using cell confluency mask in IncuCyte Zoom software (version 20158A).

### Quantification and statistical analysis

Statistical significance was assessed by comparing two datasets using a *Student’s t* test or by analyzing more than two datasets with one-way ANOVA using Prism 9. The number of trials is specified in the legends where appropriate. Statistical significance is generally denoted as follows: *, *p* < 0.05, **, *p* < 0.01, ***, *p* < 0.001, ****, *p* < 0.0001, and n.s., not significant. Western blot quantification was performed using FIJI with the “Gels” function, based on data from at least three independent experiments unless otherwise specified. All bar graphs represent the mean ± SD, accompanied by individual data points.

## AUTHOR CONTRIBUTIONS

J.P. conceived the project. O.K. and J.P. performed most of the experiments. B.G. and V.M.V. characterized the Rab7-KO cells, J.J. performed ML-SA5 experiments, and A.Q.B. and T.K. measured breast cancer cell migration and growth. O.K., B.M., A.Q.B., and J.P. analyzed the data. J.P. wrote the manuscript. All the authors edited the manuscript.

## ACKNOWLEDGMENTS

We thank J. Bonifacino for kind gift of kinesin1/3-KO cells and M. Rosas Lemus, M. Mandell, A. Kell, M. Ozbun, and J. Rajaiya for the insightful discussion. Schematic cartoons were generated by using BioRender. This research made use of the CellInsight Microscope Platform and confocal microscope from Autophagy, Inflammation, and Metabolism Center of Biomedical Research Excellence, funded by NIGMS, NIH (P20GM121176). The funding support was from NIH/NIGMS T32GM144834 and R35GM147419.

## SUPPLEMENTAL INFORMATION

Supplemental Information includes three figures and two tables.

## REFERENCES

1. J. Pu, C.M. Guardia, T. Keren-Kaplan, J.S. Bonifacino, Mechanisms and functions of lysosome positioning, J Cell Sci 129(23) (2016) 4329–4339.

2. A. Ballabio, J.S. Bonifacino, Lysosomes as dynamic regulators of cell and organismal homeostasis, Nat Rev Mol Cell Biol 21(2) (2020) 101–118.

3. M.L. Jongsma, I. Berlin, R.H. Wijdeven, L. Janssen, G.M. Janssen, M.A. Garstka, H. Janssen, M. Mensink, P.A. van Veelen, R.M. Spaapen, J. Neefjes, An ER-Associated Pathway Defines Endosomal Architecture for Controlled Cargo Transport, Cell 166(1) (2016) 152–66.

4. G.P. Starling, Y.Y. Yip, A. Sanger, P.E. Morton, E.R. Eden, M.P. Dodding, Folliculin directs the formation of a Rab34-RILP complex to control the nutrient-dependent dynamic distribution of lysosomes, EMBO Rep 17(6) (2016) 823–41.

5. M. Encarnacao, L. Espada, C. Escrevente, D. Mateus, J. Ramalho, X. Michelet, I. Santarino, V.W. Hsu, M.B. Brenner, D.C. Barral, O.V. Vieira, A Rab3a-dependent complex essential for lysosome positioning and plasma membrane repair, J Cell Biol 213(6) (2016) 631–40.

6. J.S. Bonifacino, J. Neefjes, Moving and positioning the endolysosomal system, Curr Opin Cell Biol 47 (2017) 1–8.

7. B. Cabukusta, J. Neefjes, Mechanisms of lysosomal positioning and movement, Traffic 19(10) (2018) 761–769.

8. T. Nakata, N. Hirokawa, Point mutation of adenosine triphosphate-binding motif generated rigor kinesin that selectively blocks anterograde lysosome membrane transport, J Cell Biol 131(4) (1995) 1039–53.

9. M. Matsushita, S. Tanaka, N. Nakamura, H. Inoue, H. Kanazawa, A novel kinesin-like protein, KIF1Bbeta3 is involved in the movement of lysosomes to the cell periphery in non-neuronal cells, Traffic 5(3) (2004) 140–51.

10. C.M. Guardia, G.G. Farias, R. Jia, J. Pu, J.S. Bonifacino, BORC Functions Upstream of Kinesins 1 and 3 to Coordinate Regional Movement of Lysosomes along Different Microtubule Tracks, Cell Rep 17(8) (2016) 1950–1961.

11. J.K. Burkhardt, C.J. Echeverri, T. Nilsson, R.B. Vallee, Overexpression of the dynamitin (p50) subunit of the dynactin complex disrupts dynein-dependent maintenance of membrane organelle distribution, J Cell Biol 139(2) (1997) 469–84.

12. A. Harada, Y. Takei, Y. Kanai, Y. Tanaka, S. Nonaka, N. Hirokawa, Golgi vesiculation and lysosome dispersion in cells lacking cytoplasmic dynein, J Cell Biol 141(1) (1998) 51–9.

13. C. Rosa-Ferreira, S. Munro, Arl8 and SKIP act together to link lysosomes to kinesin-1, Dev Cell 21(6) (2011) 1171–8.

14. J. Pu, C. Schindler, R. Jia, M. Jarnik, P. Backlund, J.S. Bonifacino, BORC, a multisubunit complex that regulates lysosome positioning, Developmental cell 33(2) (2015) 176–188.

15. T. Keren-Kaplan, J.S. Bonifacino, ARL8 Relieves SKIP Autoinhibition to Enable Coupling of Lysosomes to Kinesin-1, Curr Biol 31(3) (2021) 540–554 e5.

16. T. Keren-Kaplan, A. Saric, S. Ghosh, C.D. Williamson, R. Jia, Y. Li, J.S. Bonifacino, RUFY3 and RUFY4 are ARL8 effectors that promote coupling of endolysosomes to dynein-dynactin, Nat Commun 13(1) (2022) 1506.

17. G. Kumar, P. Chawla, N. Dhiman, S. Chadha, S. Sharma, K. Sethi, M. Sharma, A. Tuli, RUFY3 links Arl8b and JIP4-Dynein complex to regulate lysosome size and positioning, Nat Commun 13(1) (2022) 1540.

18. C. Raiborg, E.M. Wenzel, N.M. Pedersen, H. Olsvik, K.O. Schink, S.W. Schultz, M. Vietri, V. Nisi, C. Bucci, A. Brech, T. Johansen, H. Stenmark, Repeated ER-endosome contacts promote endosome translocation and neurite outgrowth, Nature 520(7546) (2015) 234-8.

19. Z. Hong, N.M. Pedersen, L. Wang, M.L. Torgersen, H. Stenmark, C. Raiborg, PtdIns3P controls mTORC1 signaling through lysosomal positioning, J Cell Biol 216(12) (2017) 4217–4233.

20. G. Cantalupo, P. Alifano, V. Roberti, C.B. Bruni, C. Bucci, Rab-interacting lysosomal protein (RILP): the Rab7 effector required for transport to lysosomes, EMBO J 20(4) (2001) 683–93.

21. I. Jordens, M. Fernandez-Borja, M. Marsman, S. Dusseljee, L. Janssen, J. Calafat, H. Janssen, R. Wubbolts, J. Neefjes, The Rab7 effector protein RILP controls lysosomal transport by inducing the recruitment of dynein-dynactin motors, Curr Biol 11(21) (2001) 1680–5.

22. R. Marwaha, S.B. Arya, D. Jagga, H. Kaur, A. Tuli, M. Sharma, The Rab7 effector PLEKHM1 binds Arl8b to promote cargo traffic to lysosomes, J Cell Biol 216(4) (2017) 1051–1070.

23. M.L. Jongsma, J. Bakker, B. Cabukusta, N. Liv, D. van Elsland, J. Fermie, J.L. Akkermans, C. Kuijl, S.Y. van der Zanden, L. Janssen, D. Hoogzaad, R. van der Kant, R.H. Wijdeven, J. Klumperman, I. Berlin, J. Neefjes, SKIP-HOPS recruits TBC1D15 for a Rab7-to-Arl8b identity switch to control late endosome transport, EMBO J 39(6) (2020) e102301.

24. V.I. Korolchuk, S. Saiki, M. Lichtenberg, F.H. Siddiqi, E.A. Roberts, S. Imarisio, L. Jahreiss, S. Sarkar, M. Futter, F.M. Menzies, C.J. O’Kane, V. Deretic, D.C. Rubinsztein, Lysosomal positioning coordinates cellular nutrient responses, Nat Cell Biol 13(4) (2011) 453–60.

25. M. Xu, X.X. Li, J. Xiong, M. Xia, E. Gulbins, Y. Zhang, P.L. Li, Regulation of autophagic flux by dynein-mediated autophagosomes trafficking in mouse coronary arterial myocytes, Biochim Biophys Acta 1833(12) (2013) 3228–3236.

26. R. Jia, J.S. Bonifacino, Lysosome Positioning Influences mTORC2 and AKT Signaling, Mol Cell 75(1) (2019) 26–38 e3.

27. P.A. Filipek, M.E.G. de Araujo, G.F. Vogel, C.H. De Smet, D. Eberharter, M. Rebsamen, E.L. Rudashevskaya, L. Kremser, T. Yordanov, P. Tschaikner, B.G. Furnrohr, S. Lechner, T. Dunzendorfer-Matt, K. Scheffzek, K.L. Bennett, G. Superti-Furga, H.H. Lindner, T. Stasyk, L.A. Huber, LAMTOR/Ragulator is a negative regulator of Arl8b- and BORC-dependent late endosomal positioning, J Cell Biol 216(12) (2017) 4199–4215.

28. J. Pu, T. Keren-Kaplan, J.S. Bonifacino, A Ragulator-BORC interaction controls lysosome positioning in response to amino acid availability, J Cell Biol 216(12) (2017) 4183–4197.

29. Z.N. Ling, Y.F. Jiang, J.N. Ru, J.H. Lu, B. Ding, J. Wu, Amino acid metabolism in health and disease, Signal Transduct Target Ther 8(1) (2023) 345.

30. M. Platten, E.A.A. Nollen, U.F. Rohrig, F. Fallarino, C.A. Opitz, Tryptophan metabolism as a common therapeutic target in cancer, neurodegeneration and beyond, Nat Rev Drug Discov 18(5) (2019) 379–401.

31. D.A. Guertin, D.M. Sabatini, Defining the role of mTOR in cancer, Cancer Cell 12(1) (2007) 9–22.

32. S. Hamalisto, M. Jaattela, Lysosomes in cancer-living on the edge (of the cell), Curr Opin Cell Biol 39 (2016) 69–76.

33. D. Mossmann, S. Park, M.N. Hall, mTOR signalling and cellular metabolism are mutual determinants in cancer, Nat Rev Cancer 18(12) (2018) 744–757.

34. P. Nicklin, P. Bergman, B. Zhang, E. Triantafellow, H. Wang, B. Nyfeler, H. Yang, M. Hild, C. Kung, C. Wilson, V.E. Myer, J.P. MacKeigan, J.A. Porter, Y.K. Wang, L.C. Cantley, P.M. Finan, L.O. Murphy, Bidirectional transport of amino acids regulates mTOR and autophagy, Cell 136(3) (2009) 521–34.

35. A. Elorza, I. Soro-Arnaiz, F. Melendez-Rodriguez, V. Rodriguez-Vaello, G. Marsboom, G. de Carcer, B. Acosta-Iborra, L. Albacete-Albacete, A. Ordonez, L. Serrano-Oviedo, J.M. Gimenez-Bachs, A. Vara-Vega, A. Salinas, R. Sanchez-Prieto, R. Martin del Rio, F. Sanchez-Madrid, M. Malumbres, M.O. Landazuri, J. Aragones, HIF2alpha acts as an mTORC1 activator through the amino acid carrier SLC7A5, Mol Cell 48(5) (2012) 681–91.

36. A.K. Najumudeen, F. Ceteci, S.K. Fey, G. Hamm, R.T. Steven, H. Hall, C.J. Nikula, A. Dexter, T. Murta, A.M. Race, D. Sumpton, N. Vlahov, D.M. Gay, J.R.P. Knight, R. Jackstadt, J.D.G. Leach, R.A. Ridgway, E.R. Johnson, C. Nixon, A. Hedley, K. Gilroy, W. Clark, S.B. Malla, P.D. Dunne, G. Rodriguez-Blanco, S.E. Critchlow, A. Mrowinska, G. Malviya, D. Solovyev, G. Brown, D.Y. Lewis, G.M. Mackay, D. Strathdee, S. Tardito, E. Gottlieb, C.R.G.C. Consortium, Z. Takats, S.T. Barry, R.J.A. Goodwin, J. Bunch, M. Bushell, A.D. Campbell, O.J. Sansom, The amino acid transporter SLC7A5 is required for efficient growth of KRAS-mutant colorectal cancer, Nat Genet 53(1) (2021) 16–26.

37. Y. Kanai, Amino acid transporter LAT1 (SLC7A5) as a molecular target for cancer diagnosis and therapeutics, Pharmacol Ther 230 (2022) 107964.

38. D.C. Barral, L. Staiano, C. Guimas Almeida, D.F. Cutler, E.R. Eden, C.E. Futter, A. Galione, A.R.A. Marques, D.L. Medina, G. Napolitano, C. Settembre, O.V. Vieira, J. Aerts, P. Atakpa-Adaji, G. Bruno, A. Capuozzo, E. De Leonibus, C. Di Malta, C. Escrevente, A. Esposito, P. Grumati, M.J. Hall, R.O. Teodoro, S.S. Lopes, J.P. Luzio, J. Monfregola, S. Montefusco, F.M. Platt, R. Polishchuck, M. De Risi, I. Sambri, C. Soldati, M.C. Seabra, Current methods to analyze lysosome morphology, positioning, motility and function, Traffic 23(5) (2022) 238–269.

39. C.D. Williamson, C.M. Guardia, R. De Pace, J.S. Bonifacino, A. Saric, Measurement of Lysosome Positioning by Shell Analysis and Line Scan, Methods Mol Biol 2473 (2022) 285–306.

40. R. Kamentseva, V. Kosheverova, M. Kharchenko, M. Zlobina, A. Salova, T. Belyaeva, N. Nikolsky, E. Kornilova, Functional cycle of EEA1-positive early endosome: Direct evidence for pre-existing compartment of degradative pathway, PLoS One 15(5) (2020) e0232532.

41. R. Jia, C.M. Guardia, J. Pu, Y. Chen, J.S. Bonifacino, BORC coordinates encounter and fusion of lysosomes with autophagosomes, Autophagy 13(10) (2017) 1648–1663.

42. L. Cong, F.A. Ran, D. Cox, S. Lin, R. Barretto, N. Habib, P.D. Hsu, X. Wu, W. Jiang, L.A. Marraffini, F. Zhang, Multiplex genome engineering using CRISPR/Cas systems, Science 339(6121) (2013) 819-23.

43. D.J.H. van den Boomen, A. Sienkiewicz, I. Berlin, M.L.M. Jongsma, D.M. van Elsland, J.P. Luzio, J.J.C. Neefjes, P.J. Lehner, A trimeric Rab7 GEF controls NPC1-dependent lysosomal cholesterol export, Nat Commun 11(1) (2020) 5559.

44. N. Rocha, C. Kuijl, R. van der Kant, L. Janssen, D. Houben, H. Janssen, W. Zwart, J. Neefjes, Cholesterol sensor ORP1L contacts the ER protein VAP to control Rab7-RILP-p150 Glued and late endosome positioning, J Cell Biol 185(7) (2009) 1209–25.

45. R. Willett, J.A. Martina, J.P. Zewe, R. Wills, G.R.V. Hammond, R. Puertollano, TFEB regulates lysosomal positioning by modulating TMEM55B expression and JIP4 recruitment to lysosomes, Nat Commun 8(1) (2017) 1580.

46. N. Kelkar, C.L. Standen, R.J. Davis, Role of the JIP4 scaffold protein in the regulation of mitogen-activated protein kinase signaling pathways, Mol Cell Biol 25(7) (2005) 2733–43.

47. X. Li, N. Rydzewski, A. Hider, X. Zhang, J. Yang, W. Wang, Q. Gao, X. Cheng, H. Xu, A molecular mechanism to regulate lysosome motility for lysosome positioning and tubulation, Nat Cell Biol 18(4) (2016) 404–17.

48. J.L. Jewell, Y.C. Kim, R.C. Russell, F.X. Yu, H.W. Park, S.W. Plouffe, V.S. Tagliabracci, K.L. Guan, Metabolism. Differential regulation of mTORC1 by leucine and glutamine, Science 347(6218) (2015) 194-8.

49. D. Meng, Q. Yang, H. Wang, C.H. Melick, R. Navlani, A.R. Frank, J.L. Jewell, Glutamine and asparagine activate mTORC1 independently of Rag GTPases, J Biol Chem 295(10) (2020) 2890–2899.

50. M. Rebsamen, L. Pochini, T. Stasyk, M.E. de Araujo, M. Galluccio, R.K. Kandasamy, B. Snijder, A. Fauster, E.L. Rudashevskaya, M. Bruckner, S. Scorzoni, P.A. Filipek, K.V. Huber, J.W. Bigenzahn, L.X. Heinz, C. Kraft, K.L. Bennett, C. Indiveri, L.A. Huber, G. Superti-Furga, SLC38A9 is a component of the lysosomal amino acid sensing machinery that controls mTORC1, Nature 519(7544) (2015) 477-81.

51. S. Wang, Z.Y. Tsun, R.L. Wolfson, K. Shen, G.A. Wyant, M.E. Plovanich, E.D. Yuan, T.D. Jones, L. Chantranupong, W. Comb, T. Wang, L. Bar-Peled, R. Zoncu, C. Straub, C. Kim, J. Park, B.L. Sabatini, D.M. Sabatini, Metabolism. Lysosomal amino acid transporter SLC38A9 signals arginine sufficiency to mTORC1, Science 347(6218) (2015) 188-94.

52. G.A. Wyant, M. Abu-Remaileh, R.L. Wolfson, W.W. Chen, E. Freinkman, L.V. Danai, M.G. Vander Heiden, D.M. Sabatini, mTORC1 Activator SLC38A9 Is Required to Efflux Essential Amino Acids from Lysosomes and Use Protein as a Nutrient, Cell 171(3) (2017) 642–654 e12.

53. M.A. Brandan, H.A. Perez, A. Disalvo, A.F.M. de Los, Interaction of L-phenylalanine with carbonyl groups in mixed lipid membranes, Biochim Biophys Acta Biomembr 1866(5) (2024) 184328.

54. R.L. Wolfson, L. Chantranupong, R.A. Saxton, K. Shen, S.M. Scaria, J.R. Cantor, D.M. Sabatini, Sestrin2 is a leucine sensor for the mTORC1 pathway, Science 351(6268) (2016) 43-8.

55. T.H. Kim, N.K. Gill, K.D. Nyberg, A.V. Nguyen, S.V. Hohlbauch, N.A. Geisse, C.J. Nowell, E.K. Sloan, A.C. Rowat, Cancer cells become less deformable and more invasive with activation of beta-adrenergic signaling, J Cell Sci 129(24) (2016) 4563–4575.

