## Supplemental materials for "Synergistic Role of Amino Acids in Enhancing mTOR Activation Through Lysosome Positioning"

### SUPPLEMENTAL INFORMATION

#### Supplemental figure legends

**Fig. S1. Amino acids regulate endosome and lysosome positioning**

**A.** Monkey kidney fibroblast-like cells COS-7 and human osteosarcoma U2OS cells were starved in amino acid-free DMEM media (with dialyzed serum and glucose) for 1 h, subjected to immunofluorescence with the indicated antibodies, and imaged by confocal microscopy. Yellow doted lines were manually added to indicate cell boundaries. **B.** HeLa cells were incubated in complete media, amino acid- and serum-free DMEM (Starvation) for 1.5 h, or 2 mM phenylalanine in amino acid- and serum-free media for 30 minutes following 1 h starvation. The indicated antibodies were applied in immunofluorescence, and images were collected by confocal microscopy. Scale bars, 5 μm.

**Fig. S2. The effect of amino acids on lysosome retrograde transport**

**A.** Wildtype (WT) and Rab7-KO HeLa cells were fixed and co-stained with filipin (for free cholesterol) and the indicated antibodies in detergent-free saline buffer. Confocal microscopy was performed for imaging. **B**. HeLa cells were pretreated with DMSO or 0.5 $\mu$M TRPML1 agonist ML-SA5 for 1 h, starved in amino acid- and serum-free DMEM with DMSO or ML-SA5 for 1 h, and refed with 2 mM individual indicated amino acids accompanied with DMSO or ML-SA5 for 30 min. Cells were subjected to immunofluorescence and analyzed for lysosome positioning as described in Methods. $p$ values were determined using *Student's t* test (vs. the corresponding point in the control group). ns, not significant. Scale bars, 5 μm.

**Fig. S3 Synergic effect of amino acids on mTOR activation**

**A.** HeLa cells were starved in amino acid- and serum-free DMEM for 1 h and refed with 2 mM individual or combined indicated amino acids for 30 min. Cells were subjected to immunoblotting with the indicated antibodies. The ratio of p-S6K to total S6K for each treatment was normalized to the values in the starvation group. Individual amino acid treatments were summed (labeled as "+") and compared to the corresponding values from the combination of the two amino acids (labeled as "&"). Paired values from three independent experiments were shown with lines connecting the data points. B. Wildtype (WT) and the indicated KO cells (SLC: SLC38A9. Kin: kinesin1/3) were treated as in A with combined amino acids. Paired values from two independent experiments were shown with lines connecting the data points. **C.** HeLa cells were starved in amino acid- and serum-free DMEM for 1 h and refed with 2 mM individual or combined indicated amino acids for 30 min. Cells were subjected to immunofluorescence and analysis of lysosome positioning as described in Methods. *p* values were determined using one-way ANOVA (vs. Starvation group) or *Student's t* test (indicated with lines). **, $p<0.01$ ***, $p<0.001$ ****, $p<0.0001$. ns, not significant.

**Supplemental tables**

**Table 1. Amino acid concentrations in DMEM and adults' blood**

| Amino Acid | Sigma Cat# | Stock concentration (mM) | Concentration in DMEM (mM) | Concentration in blood (mM) |
|---|---|---|---|---|
| L-Glutamine | A2916801 (ThermoFisher) | 200 | 4 | 0.4-0.8 |
| Glycine | G7126 | 200 | 0.4 | 0.12-0.55 |
| L-Arginine | 11009 | 200 | 0.4 | 0.02-0.14 |
| L-Cystine dihydrochloride | C6727-25G | (Working concentration) | 0.2 | 0.0008-0.028 |
| L-Histidine | H8000 | 200 | 0.2 | 0.057-0.114 |
| L-Isoleucine | W527602 | 200 | 0.8 | 0.038-0.13 |
| L-Leucine | 61819 | 100 | 0.8 | 0.074-0.196 |
| L-Lysine hydrochloride | L8662 | 200 | 0.8 | 0.12-0.318 |
| L-Methionine | 64319 | 200 | 0.2 | 0.014-0.148 |
| L-Phenylalanine | P2126 | 100 | 0.4 | 0.035-0.085 |
| L-Serine | S4500 | 200 | 0.4 | 0.06-0.172 |
| L-Threonine | T8625 | 200 | 0.8 | 0.073-0.216 |
| L-Tryptophan | T3300 | 50 | 0.08 | 0.031-0.083 |
| L-Tyrosine disodium salt dihydrate | RES3156T-A7 | 200 | 0.4 | 0.03-0.12 |
| L-Valine | 94619 | 200 | 0.8 | 0.146-0.37 |

**Table. S2 Antibodies used in this study**

| Antibody | Source | Product Number | Dilution | Applications |
|---|---|---|---|---|
| Actin | BD Bioscience | 612657 | 1:10,000 | WB |
| Calnexin | Cell Signaling Technologies | 2679s | 1:8,000 | WB |
| COXIV | Cell Signaling Technologies | 11967S | 1:200 | IF |
| EEA1 | Cell Signaling Technologies | 3288 | 1:100 | IF |
| JIP4 | Cell Signaling Technologies | 5519s | 1:100, 1:1,000 | IF, WB |
| LAMP1 | Cell Signaling Technologies | 9091s | 1:1,000 | IF |
| LAMP2 | Santa Cruz | SC-18822 | 1:500 | IF |
| LAMTOR4 | Cell Signaling Technologies | 13140S | 1:200 | IF |
| LC3 | MBL | PM036 | 1:200 | IF |
| PLIN2 | Santa Cruz | sc-377429 | 1:100 | IF |
| Rab5 | Cell Signaling Technologies | 3547 | 1:100 | IF |
| Rab7a | Cell Signaling Technologies | 9367 | 1:100, 1:1000 | IF, WB |
| p-S6K | Cell Signaling Technologies | 9234S | 1:1,000 | WB |
| S6K | Cell Signaling Technologies | 34475S | 1:1,000 | WB |
| TGN46 | BioRad | AHP500GT | 1:1000 | IF |

**Fig. S1**

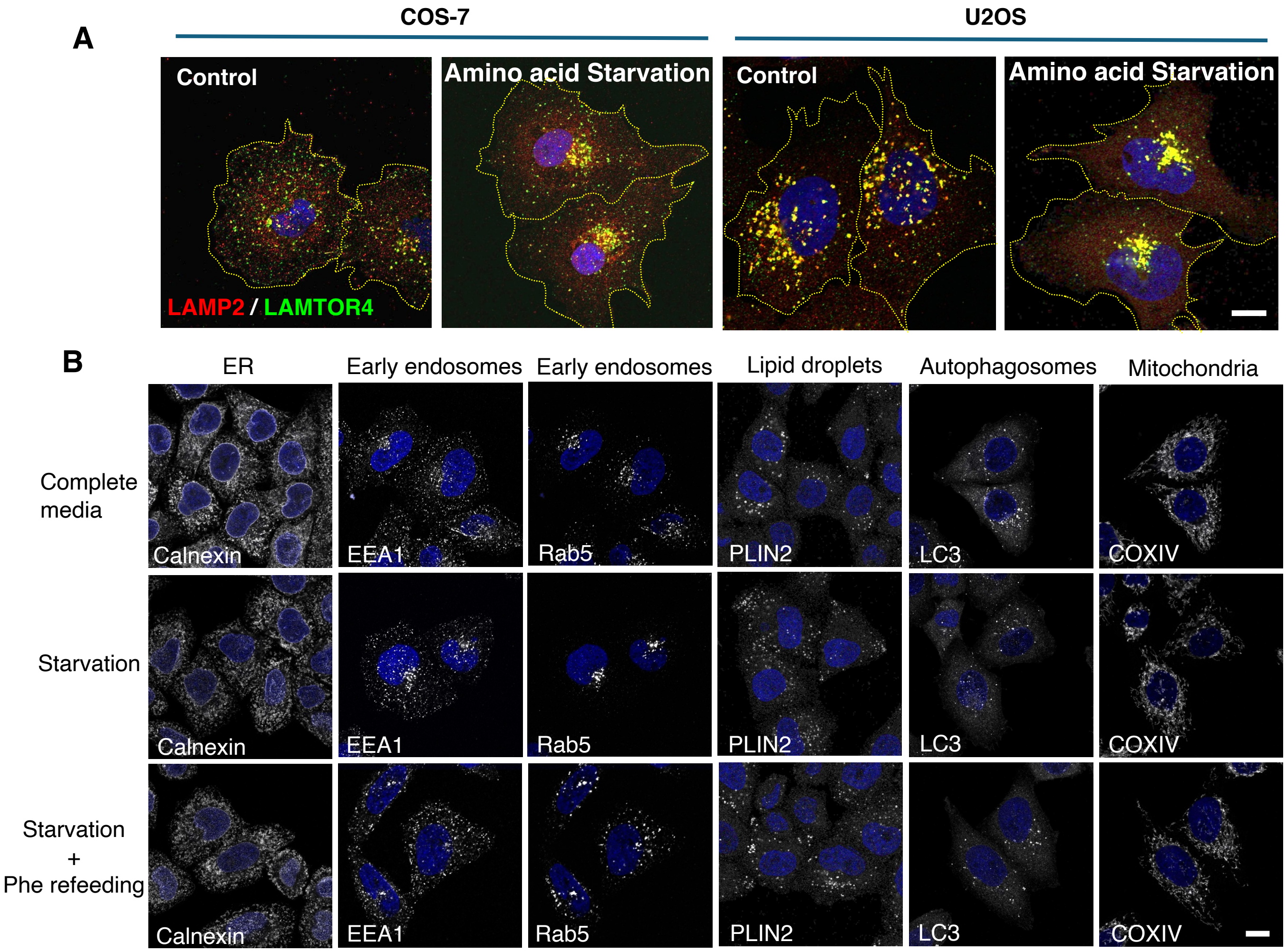
A
COS-7
U2OS
Control
Amino acid Starvation
Control
Amino acid Starvation
LAMP2 / LAMTOR4
B
ER
Early endosomes
Early endosomes
Lipid droplets
Autophagosomes
Mitochondria
Complete media
Starvation
Starvation + Phe refeeding
Calnexin
EEA1
Rab5
PLIN2
LC3
COXIV

### Fig. S2

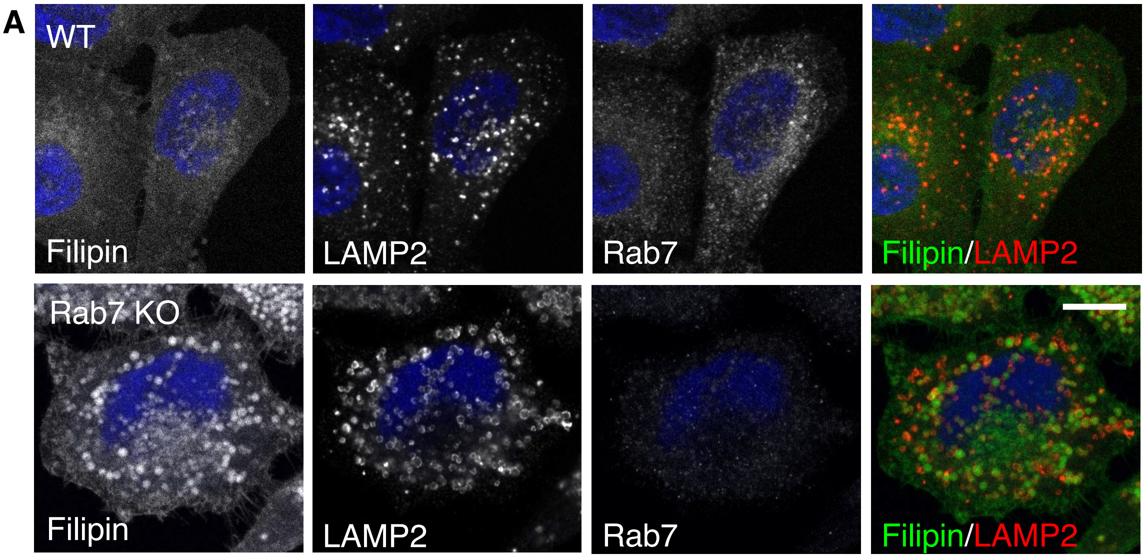

A
WT
Filipin
LAMP2
Rab7
Filipin/LAMP2
Rab7 KO
Filipin
LAMP2
Rab7
Filipin/LAMP2


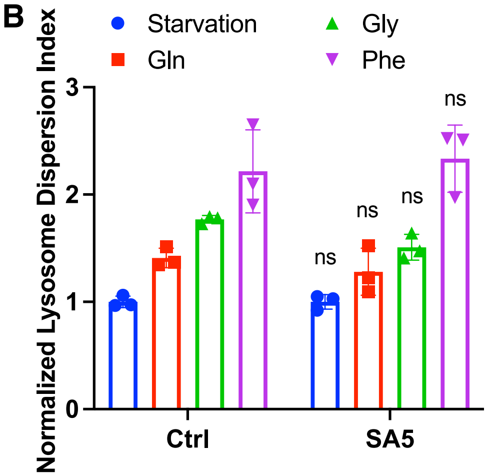

B
Starvation
Gln
Gly
Phe
Normalized Lysosome Dispersion Index
0
1
2
3
ns
ns
ns
ns
Ctrl
SA5

**Fig. S3**

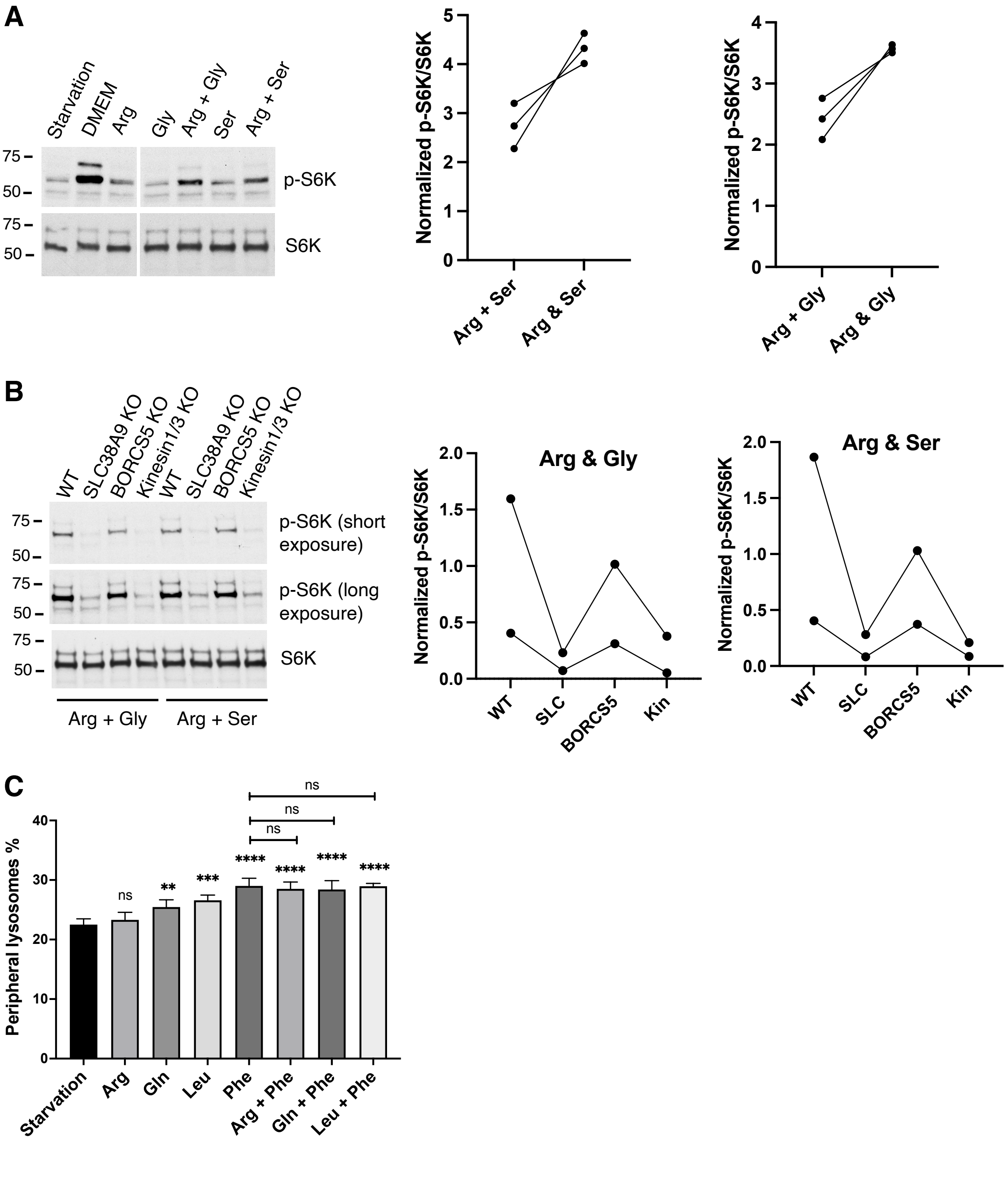
A
Starvation
DMEM
Arg
Gly
Arg + Gly
Ser
Arg + Ser
75
50
p-S6K
S6K
Normalized p-S6K/S6K
Arg + Ser
Arg & Ser
Arg + Gly
Arg & Gly
B
WT
SLC38A9 KO
BORCS5 KO
Kinesin1/3 KO
p-S6K (short exposure)
p-S6K (long exposure)
S6K
Arg + Gly
Arg + Ser
Arg & Gly
Arg & Ser
SLC
BORCS5
Kin
C
Peripheral lysosomes %
ns
**
***
****
Starvation
Arg
Gln
Leu
Phe
Arg + Phe
Gln + Phe
Leu + Phe
